# Phenformin’s impact on lifespan in *C. elegans* is resilient to environmental factors that inhibit metformin-induced longevity downstream of *skn-1/*Nrf and AMP-activated protein kinase

**DOI:** 10.1101/2023.09.21.558710

**Authors:** Sainan Li, Fasih Ahsan, Yifei Zhou, Armen Yerevanian, Alexander A. Soukas

**Affiliations:** Diabetes Unit and Center for Genomic Medicine, Massachusetts General Hospital, Harvard Medical School

## Abstract

Despite being principally prescribed to treat type 2 diabetes, biguanides, especially metformin and phenformin, have been shown to extend lifespan and healthspan in preclinical models, and to reduce the impact of aging-associated diseases such as cancer. While there have been conflicting results in studies involving rodents and humans, consistent evidence from laboratories worldwide, including our own, indicates metformin and phenformin’s ability to significantly extend lifespan in *C. elegans*. However, the pro-longevity effect of metformin can vary depending on environmental conditions. Specifically, the choice of agar from different manufacturers or batches influences metformin’s ability to extend lifespan in *C. elegans*. We traced ability of certain agar batches to interfere with metformin-prompted lifespan extension to the presence of a factor that acts directly in the worm, independently of the bacterial food source, that prevents longevity promoting effects downstream of longevity effectors *skn-1* and AMPK. In contrast, phenformin prompts robust lifespan extension in the face of environmental changes and exhibits broad positive effects in aging across genetically diverse *Caenorhabditis* species where the impact of metformin is highly variable. Thus metformin effects in aging are impacted by heretofore unappreciated environmental factors. Phenformin may represent a more robust agent with which to understand the longevity promoting mechanisms downstream of biguanides.

## Introduction

Metformin and phenformin are members of the biguanide family of compounds, a class of drugs known for its ability to lower blood glucose levels in diabetic patients. However, phenformin was withdrawn from the market in the 1970s due to increased risk of potentially fatal lactic acidosis [1]. On the other hand, metformin has emerged as the first-line choice for the treatment of type 2 diabetes. Metformin stands out for its effectiveness in countering insulin resistance, a common characteristic of type 2 diabetes, without causing significant weight gain or hypoglycemia [2]. Its long-term cardiovascular benefits have been identified through numerous studies, including the influential UK Prospective Diabetes Study (UKPDS) in 1998 [3]. The drug’s effectiveness combined with its relatively modest side effect profile has made metformin the initial therapy of choice for managing type 2 diabetes.

The potential health benefits of biguanides extend beyond their blood sugar-lowering properties. Emerging research suggests that these medications may have utility in cancer prevention and treatment, as indicated by laboratory experiments on rodents and retrospective clinical studies [4, 5]. Recent evidence suggests that metformin may have a role in reducing viral load, mitigating the cytokine storm, and improving the prognosis of patients with COVID-19 [6, 7]. Excitingly, metformin may also promote healthy aging. Several retrospective studies have reported a reduction in all-cause mortality and age-related diseases in individuals taking metformin, independent of its effect on diabetes [8–10]. Several clinical trials investigating metformin’s role in aging are currently underway or have been completed [11–15]. Supporting evidence for the these clinical trials come from studies demonstrating metformin’s ability to extend lifespan and improve healthspan in model organisms such as *C. elegans* and mice. While the evidence regarding metformin’s impact on lifespan in rodents remains controversial [16, 17], its consistent ability to promote healthspan and lifespan in *C. elegans* across different laboratories worldwide provides compelling support for further investigation of its mechanisms of action in aging and potential translation to human health [18–21].

The mechanistic targets of biguanides in aging are still being elucidated and may involve multiple overlapping pathways. For example, through inhibition of mitochondrial complex I, biguanides disrupt the electron transport chain and subsequently affect cellular energy metabolism [22–24]. This inhibition leads to the activation of adenosine monophosphate-activated protein kinase (AMPK), a cellular energy sensor that regulates various metabolic processes, including glucose uptake and fatty acid oxidation [25]. Activation of AMPK and concomitant mechanistic target of rapamycin (mTOR) inhibition have been observed in various studies and are thought to contribute to the favorable effects of biguanides [20, 26, 27]. In addition to mitochondrial complex I inhibition, mTOR inhibition and AMPK activation, many other factors have been shown to contribute to the actions of biguanides in aging. In *C. elegans* these include the activation of SKN-1/Nrf2(Nuclear factor erythroid 2-related factor 2), a transcription factor that regulates oxidative stress response, cellular detoxification processes, and defense against pathogens [20]. The nuclear pore complex (NPC) has also been implicated in biguanides’ effects in aging, although the specific mechanisms are still being studied [21].

Biguanides have also been shown to affect microbial folate and methionine metabolism in bacteria, which can indirectly influence cellular processes such as fatty acid oxidation that promote longevity in the *C. elegans* feeding on those bacteria [18, 28]. Finally, lysosomal pathways have been invoked in the cellular response to biguanides [19, 29].

A recent study utilizing cryo-electron microscopy elucidated the structural basis for biguanide binding to mammalian mitochondrial respiratory complex I [30], making concrete the ability of biguanides to directly inhibit the electron transport chain. It is important to note that the mechanisms of metformin action depend on the specific outcome being studied. For example, *aak-2,* one of two *C. elegans* AMPK alpha catalytic subunit genes, is required for the lifespan extension effect of metformin in *C. elegans*, but not for the drug’s ability to inhibit growth in the worm [20, 21]. This highlights the complexity of metformin’s mechanisms of action and the context-dependent nature of its effects.

Metformin and phenformin, being biguanide derivatives, share similarities in their mechanism of action on mitochondrial complex I. Both drugs have been observed to preferentially bind to the inactive state of complex I, which is a unique characteristic not observed with other classes of complex I inhibitors [30]. This specific binding behavior distinguishes biguanides from other compounds that target complex I. While metformin and phenformin share similarities, phenformin exhibits certain characteristics that make it more potent than metformin: phenformin, with its terminal nitrogen atom substituted by a 2-phenylethyl group, is less polar and more lipid soluble than metformin. This higher lipid solubility allows phenformin to have a greater affinity for mitochondrial membranes [31]. Additionally, unlike the hydrophobic metformin, phenformin has a much greater affinity for organic cation transporters (OCTs) and is therefore more readily transported into cells [32–34]. It is important to note that the higher potency of phenformin comes with increased risks: phenformin’s higher risk of causing lactic acidosis is also a result of its higher lipophilicity and ability to freely enter cells. However, these features likely also contribute to phenformin’s potential as a more potent anticancer and prolongevity agent when compared to metformin. Ongoing research and studies are exploring the therapeutic applications of phenformin in these areas [35–37]. For example: phenformin exhibits higher potential in the treatment of leukemia/lymphoma compared to metformin, primarily due to its superior cellular uptake by thymocytes [37]. The risks and benefits associated with phenformin in clinical applications related to aging and aging-associated diseases remain incompletely explored.

In this study, we find that compared to metformin, phenformin’s pro-longevity effect is robust to environmental variables that compromise metformin’s effects on lifespan. We find that this robustness is not attributed to differences in uptake of phenformin versus metformin, rather to an intrinsic ability of phenformin to directly promote longevity effectors in *C. elegans*. Specifically, we find that environmental factors interfere with metformin ability to promote lifespan extension downstream of its longevity effectors *skn-1* and AMPK. These same environmental factors do not compromise phenformin-mediated lifespan extension. Additionally, phenformin is also robust to genetic differences manifest in a wide range of *Caenorhabditis* species, indicating that it may provide a more optimal scenario with which to elucidate longevity-promoting mechanisms stimulated by biguanides.

## Results

### Environmental factors abrogate metformin-but not phenformin-stimulated longevity in C. elegans

Metformin, when used at a concentration of 50 mM, extends the lifespan of *C. elegans* [18–21], an observation we and others have reproduced many times in our laboratory (Figure 1A).

**Figure 1.**
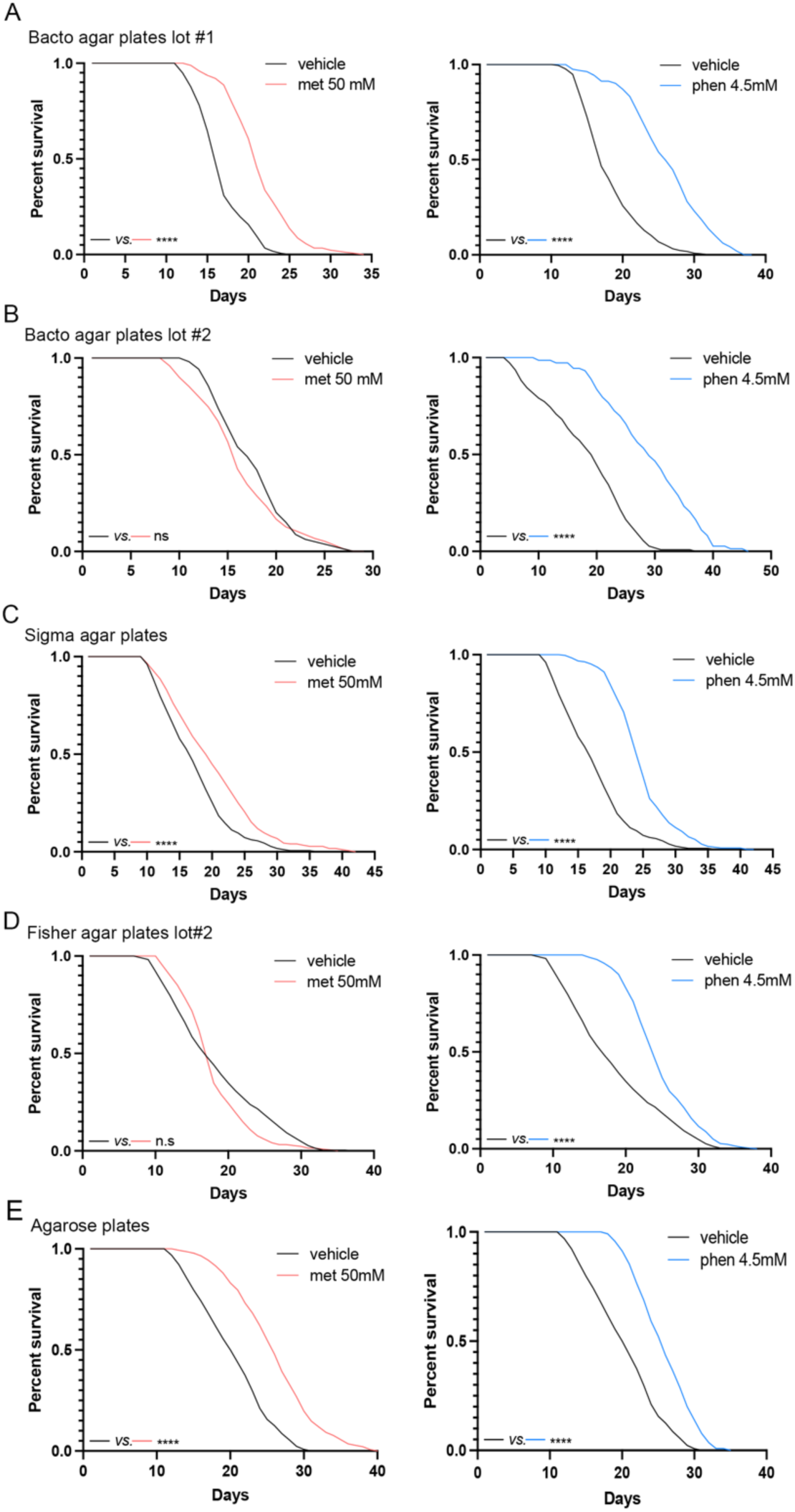
Metformin-but not phenformin-prompted lifespan extension varies with media. (A-E) Metformin extends *C. elegans* lifespan on Bacto agar plates lot #1 (A), Sigma agar plates (C), and agarose plates (E), but not in Bacto agar plates lot #2 (B) and Fisher Scientific agar plates lot #2 (D). Phenformin extends *C. elegans* lifespan in all conditions (log-rank test). Results are representative of 2–4 biological replicates (see also Table S1 for tabular results and biological replicates). ****p < 0.0001. See also Figure S1.

However, this effect suddenly and abruptly ceased (Figure 1B), an effect we traced to a different lot of agar from the same manufacturer (Bacto). To further investigate the dependence of metformin-prompted lifespan extension on agar as a growth media component, we conducted experiments using different brands and lots of agar for plate preparation. We observed that metformin’s lifespan extension effect was weakened or “blunted” in plates prepared using agar sourced from Sigma Aldrich (Figure 1C) and a specific lot of Fisher Scientific agar (Figure 1D). However, the effect remained strong in plates using a distinct lot of Fisher Scientific agar or commercially purchased *C. elegans* media plates (Figure S1A). In stark contrast, consistent and robust lifespan extension effects are seen in response to phenformin treatment of *C. elegans* across all types of agars used (Figure 1 and S1A).

We next expanded our investigation to include agarose. Interestingly, both metformin and phenformin extended lifespan on agarose plates (Figure 1E). This observation suggests that a more highly purified gelling agent, like agarose, might lack certain interfering substances present in agar, accounting for the differential outcomes.

We further investigated other potential factors that could influence metformin’s effect on lifespan, including changing the brand of metformin (Figure S1B), altering the timing of metformin administration (pre-developmental or post-developmental; Figure S1C), and modifying the culture conditions of OP50 bacteria (cold storage, shaken culture, or still culture, as detailed in methods; Figure S1D). However, none of these modifications restored the ability of metformin to extend lifespan on Bacto agar plates. Additionally, we investigated whether different media simply altered the dose response to metformin by examining the impact of higher or lower dosages of metformin (ranging from 10 mM to 100 mM) on the lifespan of worms grown on Bacto agar plates (Figure S1D and E). The results showed that regardless of the dosage used, metformin did not extend lifespan on Bacto agar plates.

Streptomycin is an antibiotic commonly employed during *C. elegans* media preparation to mitigate bacterial contamination, and its mechanism of action involves inhibiting bacterial translation. Consequently, we conducted an investigation to determine whether streptomycin effect could confound metformin’s action. Metformin does not extend the lifespan of worms when they are grown on Bacto agar plates, regardless of whether the plates were prepared with or without streptomycin (Figure 1F). In contrast, when *C. elegans* is cultured on Fisher Scientific agar plates or agarose plates, our observations demonstrate that metformin consistently extends the lifespan of the worms, irrespective of whether streptomycin was added during plate preparation (Figure S1A and G). In longevity assays, fluorodeoxyuridine (FUdR) is frequently utilized post-developmentally to inhibit *C. elegans* progeny production by interfering with DNA replication by binding to thymidylate synthase. Therefore, we conducted experiments to examine whether FUdR’s influence could account for any variations observed in metformin action. Our findings revealed that the presence or absence of FUdR did not have any effect on the impact of metformin in this context (Figure S1C). Finally, the bacterial food source does not appear to influence metformin effects, as metformin extends lifespan whether *C. elegans* are fed *E. coli* HT115 bacteria or *E. coli* OP50-1 bacteria when grown on agarose-containing plates (Figure S1H).

Based on the aforementioned findings, we can conclude that the only environmental factor that influences the variation in metformin’s effect on lifespan extension is the type of gelling agent (i.e. agar, agarose) used in *C. elegans* solid media. On the other hand, phenformin consistently extends lifespan across all experimental conditions tested (Figure 1 and S1).

### Metformin and phenformin extend lifespan additively and share the same genetic dependency as metformin or phenformin alone

In light of the observed variations in metformin’s pro-longevity effect with different media, while phenformin remains unaffected, we further investigated whether these two drugs have distinct genetic dependencies in aging. Previous studies have suggested that the energy sensor AMPK, the oxidative stress-responsive transcription factor SKN-1/Nrf2, and the nuclear pore complex (NPC) play crucial roles in metformin’s lifespan benefits in *C. elegans* [20, 21]. We find that phenformin’s lifespan extension effect is also dependent on the genetic components *aak-2*/AMPK, *skn-1*/Nrf2, as well as NPC constituents such as *npp-3* and *npp-21* (Figure S2A-C). It has been previously reported that biguanides, including metformin and phenformin, can directly interact with mitochondrial complex I [30]. In our investigation, we made a significant observation that both metformin and phenformin were unable to extend lifespan in *gas-1* mutants (Figure S2D), which are characterized by mutations in a mitochondrial complex I subunit. In aggregate, these findings indicate that metformin and phenformin share the same genetic dependencies to extend lifespan, and suggests, but does not prove, that they have similar mechanisms of action.

Curiously, when metformin and phenformin are administered together, lifespan extension is additive, surpassing lifespan extension achieved by either metformin or phenformin alone, when agarose plates are employed (Figure 2A and B). This effect is consistent regardless of whether OP50 or HT115 *E. coli* are used as the worms’ bacterial food source. Importantly, higher dosages of metformin or phenformin shorten rather than extend lifespan (Figure S1D) [18], indicating that the interaction between metformin and phenformin is likely to be complex. However, it is noteworthy that this additive lifespan benefit is not observed in conditions where metformin alone is unable to extend lifespan such as Bacto agar plates and Sigma Aldrich agar plates (Figure S3). Importantly, the additive lifespan extension effect observed when administering both metformin and phenformin together is completely dependent on the genetic factors AAK-2/AMPK, SKN-1/Nrf2, and mitochondrial complex I (Figure 2C), indicating that irrespective of the mechanism of the additivity, the lifespan extending effect of co-administered metformin and phenformin converges on the same set of downstream effectors.

**Figure 2.**
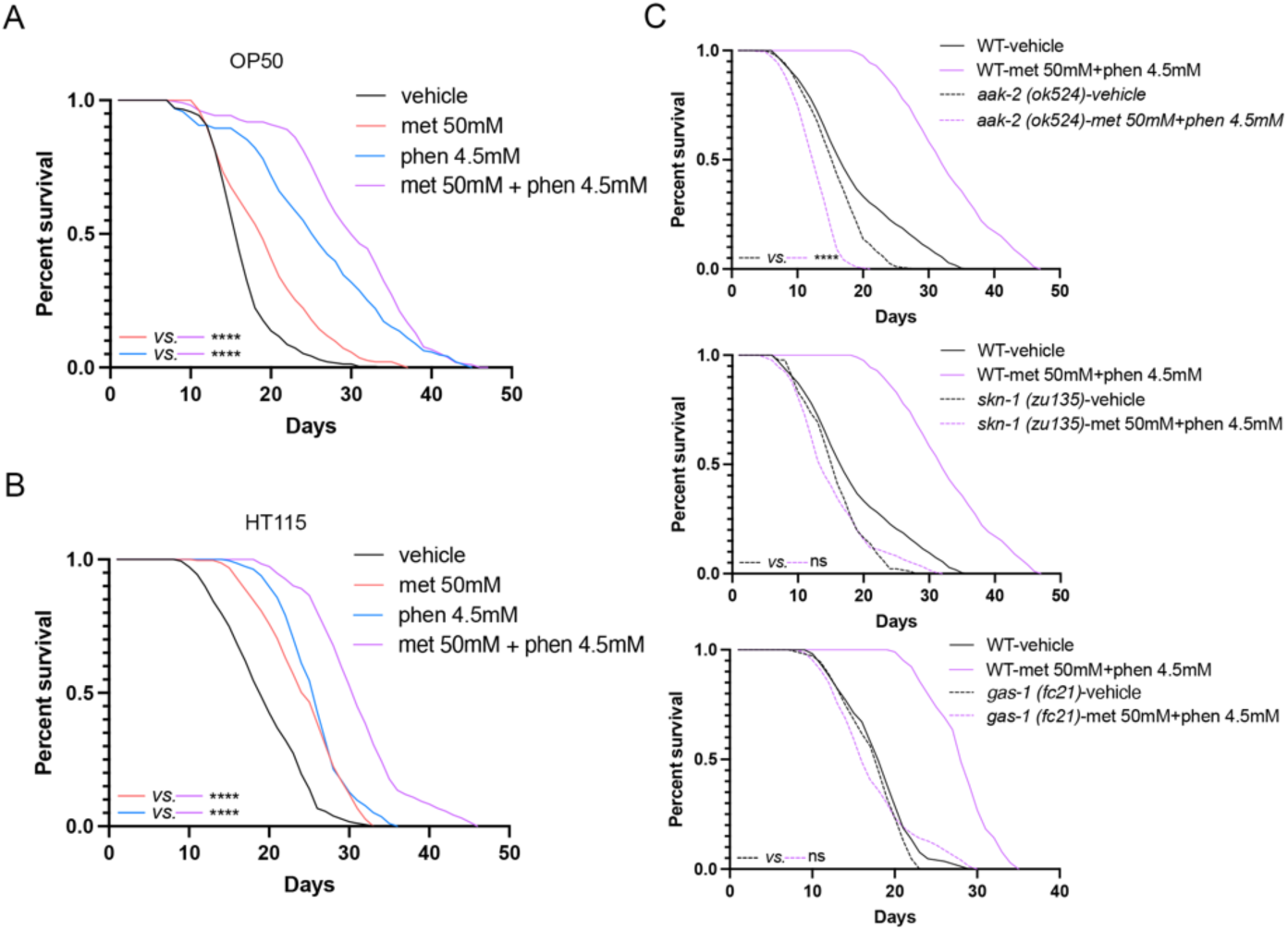
Metformin and phenformin extend lifespan additively. (A and B) When *C. elegans* is fed OP50 (A) or HT115 (B) bacteria and cultured on agarose plates, the combination of metformin and phenformin leads to an additive extension of lifespan (log-rank test). (C) On agarose plates, metformin combined with phenformin cannot extend *C. elegans* lifespan in *aak-2(ok524), skn-1(zu135)* and *gas-1(fc21)* mutants (log-rank test). Results are representative of 3–5 biological replicates (see also Table S2 for tabular data and biological replicates). ****p < 0.0001. See also Figure S2 and S3.

### The promotion of longevity in C. elegans by metformin and phenformin is not dependent on bacterial metabolism or the worm’s ability to uptake the drugs

Multiple studies have suggested that metformin may exert its hypoglycemic, anti-cancer, and lifespan extending effects by influencing the composition and metabolism of the gut microbiota [18, 38, 39]. In order to investigate whether the variation in metformin’s pro-longevity effect is mediated by shifts in bacterial metabolism on different media, we conducted lifespan analyses using paraformaldehyde (PFA)-killed bacteria, which are neither metabolically active nor capable of replication [40]. This approach allows potential confounding from alterations in bacterial metabolism to be avoided and focus given specifically to the direct impact of metformin and phenformin on *C. elegans*. On agarose plates, both metformin and phenformin extend the lifespan of worms feeding on live or PFA-killed *E. coli* OP50-1 bacteria (Figure 3A and B). This suggests that, under experimental conditions tested, that lifespan extension prompted by biguanides occurs via direct drug impact on worms, independent of bacterial metabolism. Conversely, on Bacto agar plates, phenformin can extend lifespan in worms feeding on both live and dead bacteria, whereas metformin fails to extend lifespan in both scenarios (Figure 3C).

**Figure 3.**
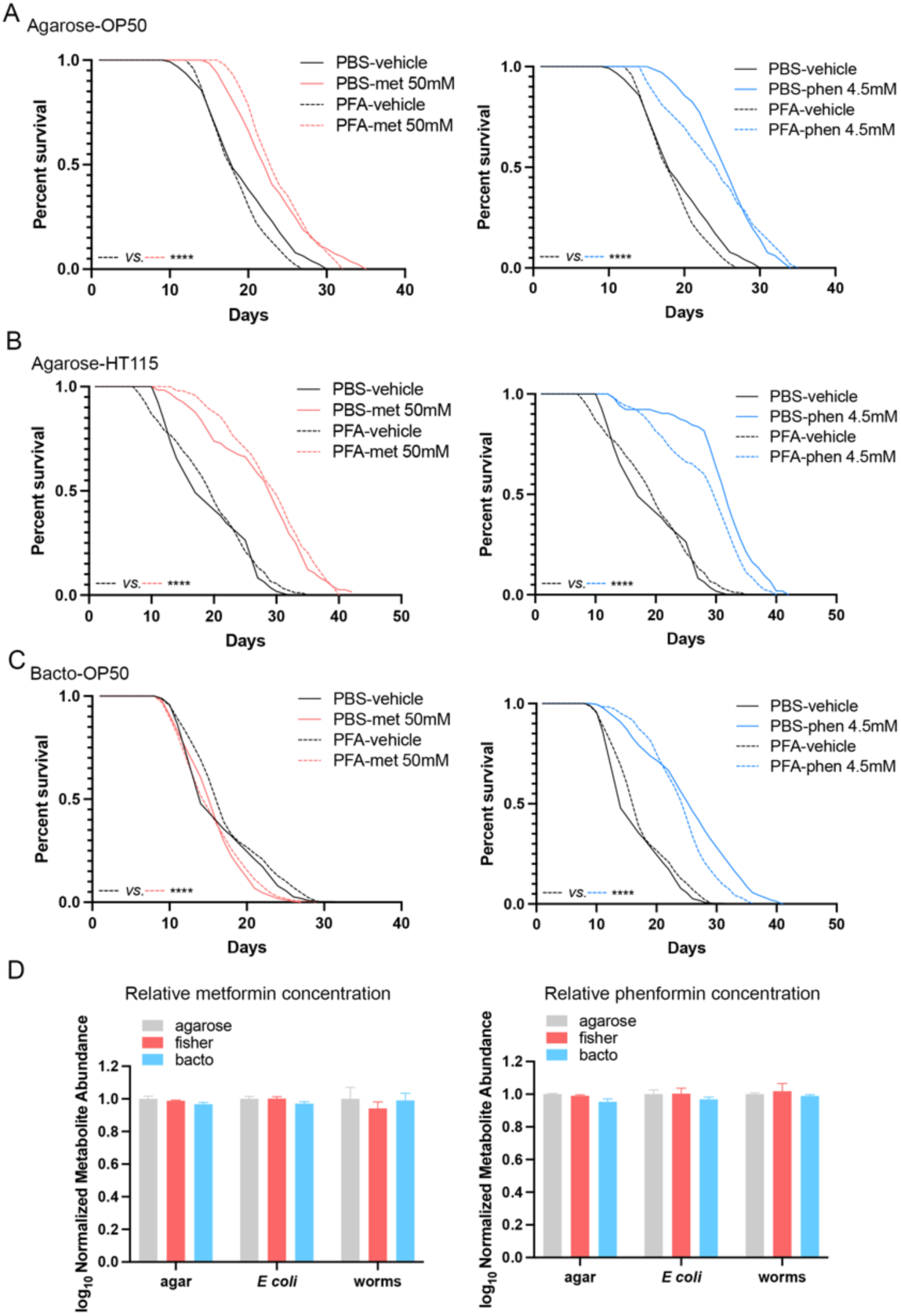
Biguanides extend the lifespan of *C. elegans* independently of bacterial metabolism. (A and B) On agarose plates, both metformin and phenformin extend the lifespan of *C. elegans* when the worms are fed with PBS-(live) or paraformaldehyde (PFA)-treated (dead) OP50 (A) or HT115 (B) bacteria (log-rank test). (C) On Bacto agar plates, phenformin, but not metformin, extends the lifespan of *C. elegans* when the worms are fed with PBS or PFA-treated OP50 bacteria (log-rank test). (D) There are no significant differences in the relative concentrations of metformin or phenformin measured by mass spectrometry in the solid media, *E. coli*, and *C. elegans* grown on agarose, Bacto agar, or Fisher agar plates. The agarose group was utilized as a control in this analysis (two-way-ANOVA). Results are representative of 2–3 biological replicates (see also Table S3 for tabular results and biological replicates). ****p < 0.0001. See also Figure S4.

These results suggest that the media component that interferes with metformin-induced lifespan extension acts directly on worms, and is not reliant on bacterial metabolism.

To further validate our conclusion that biguanides extend lifespan via direct action on *C. elegans*, we conducted experiments using UV-radiated bacteria. Exposure of bacteria to UV radiation renders them incapable of replication and most metabolic activity. Since UV irradiation leads to some bacterial lysis, worms feeding on UV-irradiated bacteria exhibit a modest degree of dietary restriction and associated lifespan extension (Figure S4). However, we observed that lower doses of phenformin (1.5 and 2 mM) are still able to extend lifespan when UV-killed bacteria are used (Figure S4B and C). Higher doses of phenformin on UV-irradiated bacteria leads to shortened lifespan (Figure S4E), emphasizing that the metabolic state of the host *C. elegans* is critical in determining the dose of biguanides capable of maximally promoting lifespan.

In order to investigate whether differences in drug uptake contribute to the variable lifespan outcomes of metformin on different plates, we utilized mass spectrometry to measure the concentrations of biguanides in worms grown on Bacto agar, Fisher Scientific agar, and agarose plates. We discovered no significant differences in drug concentrations between worms grown on different types of solid media plates (Figure 3D). This finding provides additional support to the notion that attenuation of the lifespan extending effects of metformin in response to solid media gelling agents occurs downstream of biguanide effector pathways, not by interfering with drug uptake or metabolism.

### Media mitigates metformin-induced longevity downstream of skn-1 and AMPK

Since our data indicate that the environmental factor in agar that interferes with metformin-stimulated lifespan extension acts directly in *C. elegans*, we next sought to localize more precisely the site of interference. To begin to approach this question, we asked whether lifespan modulation that is a consequence of direct activation of AAK-2/AMPK and SKN-1/Nrf2 pathways is similarly influenced by the choice of media. Our findings reveal that AAK-2 overexpression leads to a longer lifespan compared to the wild-type strain when cultured on agarose plates (Figure 4A), which is consistent with prior reports [25]. However, no significant lifespan benefits are observed in the AAK-2 overexpression strain when cultured on Bacto agar plates. On the other hand, *skn-1* gain-of-function mutants exhibited a longer lifespan than the wild-type strain when cultured on both Bacto agar and agarose plates (Figure 4B). Notably, the lifespan extension effect was more pronounced in agarose plates (3.5 days in agarose plates compared to 2 days in Bacto agar plates). The fact that direct activation of AAK-2/AMPK and SKN-1/Nrf2 also vary with media suggests that the media-induced modulation of biguanide longevity occurs downstream of *skn-1* and AMPK, rather than more proximally affecting drug levels or interaction with its immediate targets.

**Figure 4.**
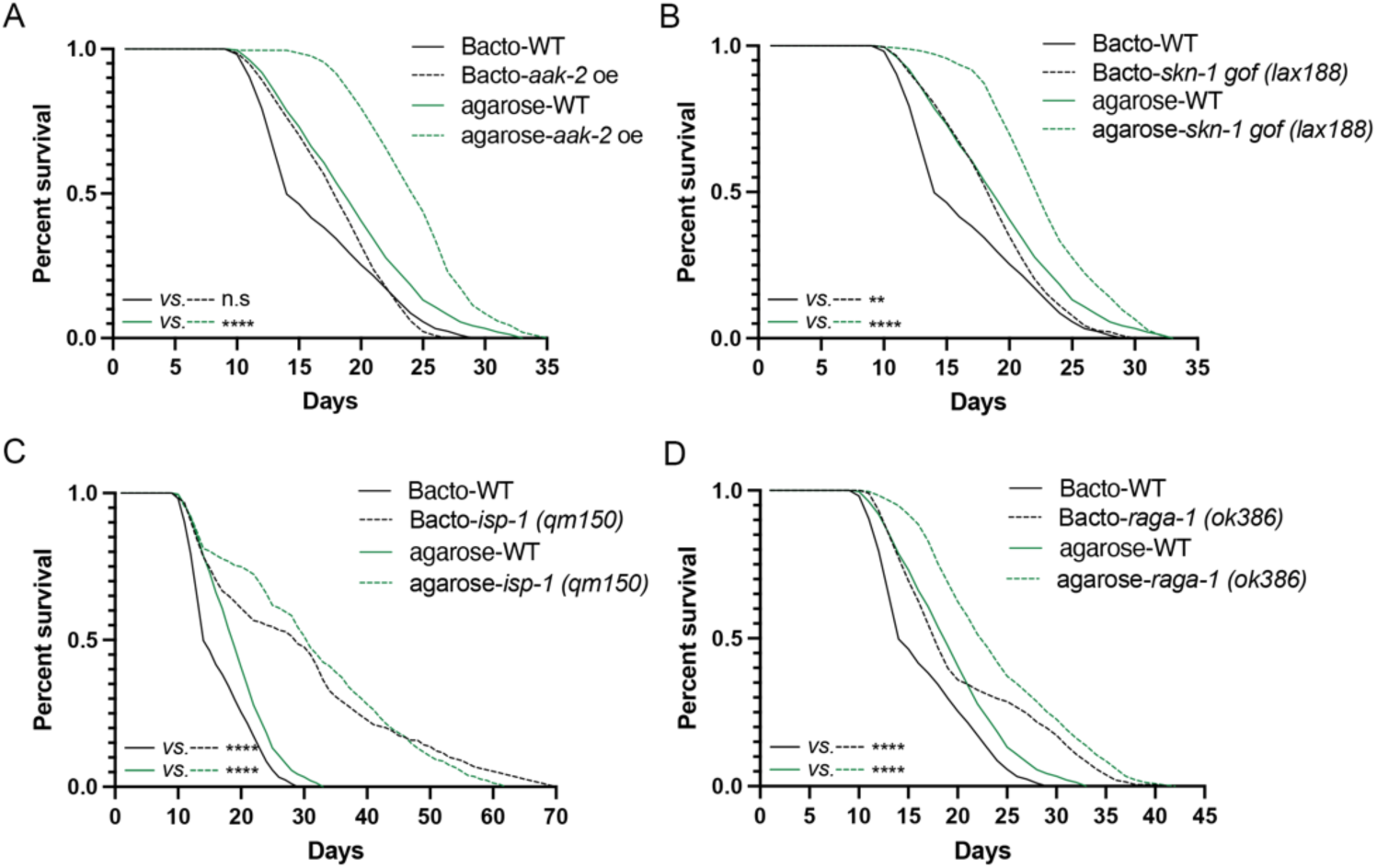
Media is a major variable impacting lifespan extension attributable to genetic activation of AMPK and *skn-1*. (A) The *aak-2* overexpression strain (*uthIs248*) outlives wild type on agarose plates but not on Bacto agar plates (log-rank test). (B) *skn-1*(*lax188*) *gain-of-function* mutants outlive wild type on agarose plates and Bacto agar plates (log-rank test). (C and D) *isp-1(qm150)* and *raga-1(ok386)* mutants outlive wild type on agarose plates and Bacto agar plates (log-rank test). Results are representative of 3 biological replicates (see also Table S4 for tabular results and biological replicates). **p < 0.01, ****p < 0.0001.

In contrast, the lifespan of mitochondrial electron transport chain deficient *isp-1* mutant and *raga-1* mutants, in which mTOR complex 1 activity is inhibited, remains unaffected by changes in the media (Figure 4C and D), indicating that the lifespan extension effects induced by mitochondria deficiency and TOR inhibition are resilient to environmental factors that interfere with metformin action in aging.

### Phenformin has a positive impact on lifespan in a broader array of Caenorhabditis species than metformin

Studies have indicated that the outcomes of metformin’s health effects in human populations are influenced by genetic variation [41]. Recent research has further suggested that metformin increases survival in certain *C. elegans* strains but not in *C. briggsae* and *C. tropicalis* strains [42]. To explore whether the attenuated pro-longevity effect of metformin on certain media applies to other genetic backgrounds, we investigated its impact on different *C. elegans* strains grown on Bacto agar plates. Our findings indicate that when using Bacto agar plates, metformin’s lifespan-extending benefits are abolished not only in the N2 strain but also in all three *C. elegans* strains tested (Figure 5A). These results indicate the principle that environmental changes determine metformin’s pro-longevity effect applies to a wider range of genetic variation. However, phenformin exhibits the ability to extend lifespan in all three *C. elegans* strains on Bacto agar plates (Figure 5A). Further unlike metformin, phenformin is able to prolong lifespan in two *C. briggsae* strains (AF16 and HK104) and two *C. tropicalis* strains (JU1373 and JU1630; Figure 5B and C). Indeed, the dosage of phenformin needed to extend lifespan varies among different nematode strains. For AF16, a dosage of 1.5mM phenformin is required. As for HK104, two different dosages are effective: 0.5mM and 1.5mM. In the case of JU1373, a dosage of 1.5mM phenformin is sufficient, while JU1630 requires a higher dosage of 4.5mM phenformin. These distinct dose-response relationships highlight the strain-specific response to phenformin and the importance of tailoring the dosage for optimal lifespan extension in each strain. These insights also shed light on the intricate interplay between genetic factors, environmental conditions, and the efficacy of metformin and phenformin in promoting longevity. It is unclear as yet whether these results suggest mechanistically distinct modes of action for metformin versus phenformin.

**Figure 5.**
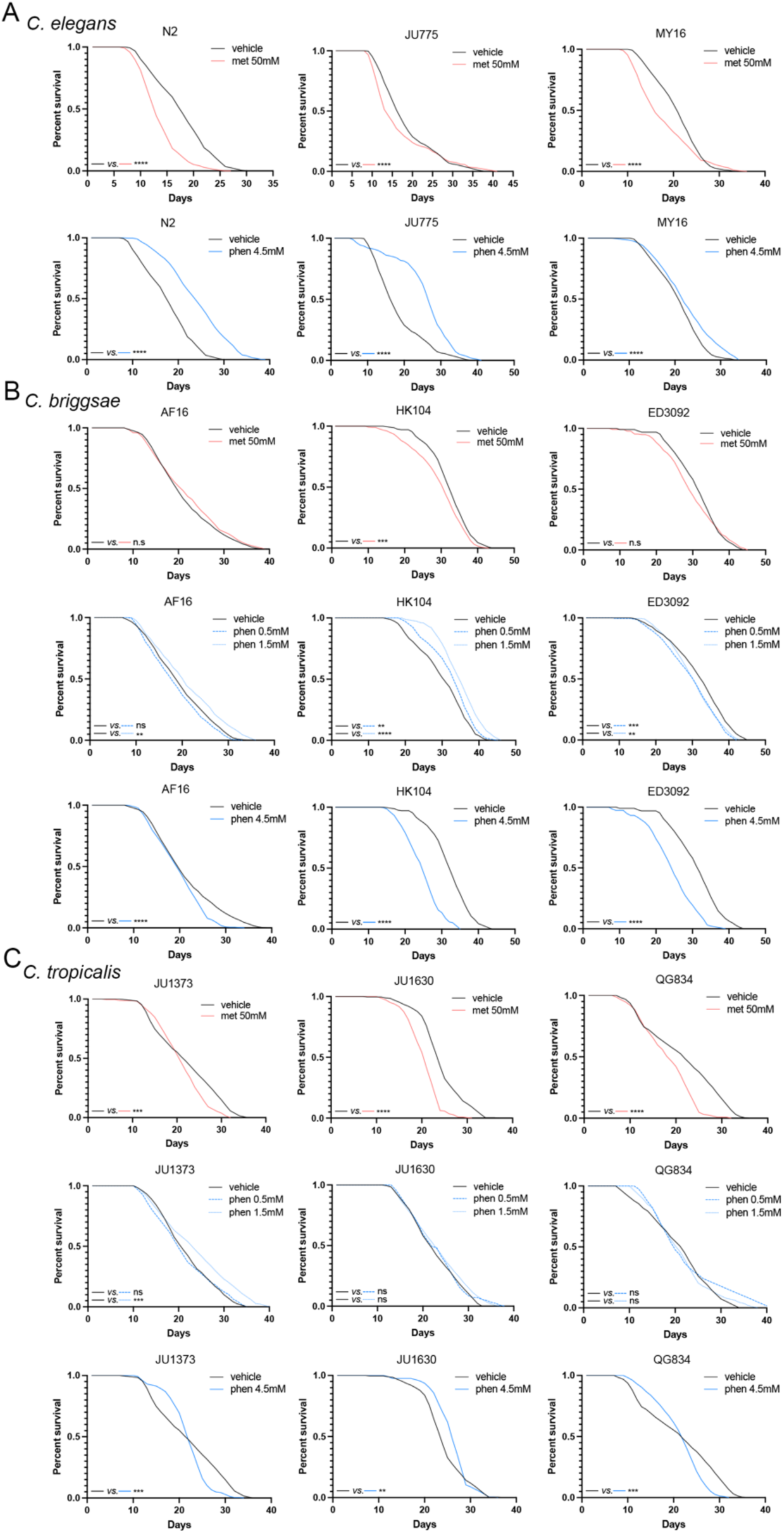
Phenformin but not metformin extend lifespan in *C. elegans, C. briggsae* and *C. tropicalis* strains using Bacto agar plates. (A) Metformin at 50 mM fails to extend lifespan, whereas phenformin at 4.5mM extends lifespan in all three *C. elegans* strains (N2, JU775, and MY16) using Bacto agar plates (log-rank test). (B) Metformin at 50 mM and phenformin at 4.5 mM fail to extend lifespan in all three *C. briggsae* strains (AF16, HK104, and ED3092) using Bacto agar plates, but phenformin at 1.5 mM extend lifespan of AF16, and phenformin at 0.5 mM and 1.5 mM extend lifespan of HK104 (log-rank test). (C) Metformin at 50 mM fail to extend lifespan in all three *C. tropicalis* strains (JU1373, JU1630, and QG834) using Bacto agar plates, but phenformin at 1.5 mM extends lifespan of JU1373, and phenformin at 4.5 mM extends lifespan of JU1630 (log-rank test). Results are representative of 3 biological replicates (see also Table S5 for tabular results and biological replicates). **p < 0.01, ***p < 0.001, ****p < 0.0001.

## Discussion

In this manuscript, we elaborate on the impact of a putative interfering substance in some but not all agar preparations that blocks the favorable effects of metformin, AAK-2 overexpression and *skn-1* gain-of-function on lifespan. Interestingly, this substance does not affect phenformin, despite both biguanides seemingly relying on identical genetic dependencies for their lifespan-promoting effects. Surprisingly, the interference from select agar preparations does not operate by modulating bacterial metabolism; instead, it directly affects *C. elegans*, likely downstream of the aforementioned longevity effectors AMPK and SKN-1. Although our results neither confirm nor refute the possibility that metformin and phenformin prompt longevity via common primary targets, in aggregate our data indicate that phenformin effect is robust to environmental variables and genetic differences manifest in diverse *Caenorhabditis* species. We propose that further investigation into the unique and shared mechanisms through which both biguanides enhance adaptive stress defenses during aging will illuminate novel strategies for promoting healthy aging. Moreover, a more in-depth understanding of the full range of environmental and genetic factors modulating the beneficial effects of these drugs in aging may allow for the optimal utilization of biguanides to improve human health.

Agar, the natural product used as a gelling agent in solid nematode growth media, is one underappreciated contributor to variable outcomes in life history traits in the worm. Agar is extracted directly from the cell wall of different species of algae, and differences in both the species of algae and the extraction conditions contribute to variations in agar’s properties. Nematode growth medium (NGM), the main media on which *C. elegans* are maintained in the laboratory [43], is a combination of agar, peptone, salts, and several other defined additives. In the data presented in this manuscript, the importance of agar in study of *C. elegans* aging emerged when lifespan results changed following a shift to a different lot of Bacto agar in the lab. As nothing else changed compared to previous conditions when metformin could extend *C. elegans* lifespan, we conclude that either a change in the process for formulation or extraction of agar, or a change in the growth conditions of the algae used to extract the agar, is responsible for the elimination of metformin effect on lifespan. Given that interference with the lifespan-promoting effect of metformin was not observed when more highly purified gelling agent agarose was used, we hypothesize that the natural product agar, which does not have a precisely defined composition, could involve any of countless component molecules. Our attempts to isolate the interfering compound from Bacto agar by methanolic extraction were unsuccessful (data not shown) suggesting that a complex approach may be required to pinpoint the exact component(s) responsible.

Having identified agar as the likely source of a putative compound that interferes with metformin-prompted lifespan extension in *C. elegans*, we next explored the potential site of action of the agar-derived substance. Numerous studies illuminate the significant impacts of bacteria, serving as a food source, on the proliferation and aging processes of *C. elegans* [44–46]. *Cabreiro et al.* reported that metformin increases *C. elegans* lifespan via impact on living bacteria, specifically by altering microbial folate and methionine metabolism [18, 28]. *Espada et al.* indicated that late-life metformin toxicity is independent of drug effects on the bacterial food source [47]. Our findings indicate that metformin increases lifespan, at least under conditions tested, via a direct impact on *C. elegans*. Further, and surprisingly, the blockade of metformin’s action by certain media (i.e. specific lots of Bacto agar) is also independent of bacterial metabolism. However, results presented here are not mutually exclusive with the possibility of bacterial metabolism playing an important role in metformin’s effect on aging, when appropriate gelling agents are used.

We demonstrate that changes in agar can lead to variation in the lifespan extending effects induced by genetic (gain-of-function or overexpression) and pharmacological (biguanides) manipulation of AMPK and SKN-1 activation. However, these effects are not universal and are likely to be multifaceted. For example, lifespan extension prompted by mTORC1 inhibition, which also depends on AMPK and *skn-1*, is not impeded by different solid media gelling agents. Furthermore, it is not simply interference with these factors, as the effect of phenformin, which also requires AMPK and *skn-1*, is observed regardless of the type of media used. We propose one potential explanation for these phenomena: even though published literature suggests that AMPK, mTORC1, and *skn-1* work in a concerted genetic pathway and highly likely at least in part in the same tissue (i.e. neurons), the mechanisms by which mTORC1 suppression, mitochondrial electron transport blockade, AMPK and SKN-1 activation promote healthy aging are likely distinct. For example, although AMPK operates genetically downstream of mTOR-mediated longevity, mutations in CRTC-1(S76A, S179A) fully suppresses AMPK-mediated longevity but do not similarly suppress *raga-1* mutant induced longevity [25, 48], suggesting complex and branching relationships of these factors.

This leads to the question of why metformin’s effects in aging vary with different media whereas phenformin’s effects are robust. One possible explanation is despite the shared genetic dependencies, more precise mechanisms by which metformin and phenformin promote longevity could be separable. Another possibility is different sites of action by metformin and phenformin, since phenformin is more lipid soluble that may have better penetrance to neurons. The third possible explanation for the segregation phenotypes is these two biguanides could be the strength of activation or inhibition on the pathway. For example, phenformin has a better affinity and is more concentrated in mitochondria. However, the fact that metformin and phenformin have an additivity effect in pro-longevity favors the first possibility. Despite prior publications suggesting that metformin and phenformin do not additively promote lifespan extension in *C. elegans* [18], our results indicate that media is a critical determinant of the outcome of this experiment.

Aging is accompanied by a gradual decline in functionality in most body systems and reduced quality of life. Among all model organisms used for fundamental study of aging, *C. elegans* stand out because of many advantages, including their short lifespan, high genetic homology (60–80%) with humans, and the availability of molecular and genetic tools [49]. However, emerging evidence suggests that environmental factors and lab-to-lab variance are critical determinants of lifespan in *C. elegans*, despite the same wild-type reference strain N2 being used. Temperature, the amount of food [50], humidity, air pressure and quality [51], use of the chemical sterilizer 5-fluoro-2′-deoxyuridine (FUdR), and even the geographic location of the laboratory have been shown to contribute to the variation observed in wild-type *C. elegans* lifespan [52]. Here we present another example of the complex impact of environment and genetics on life history traits. We emphasize the overarching conclusion that without a fuller understanding of environmental conditions that influence aging, even longevity manipulations that are well-characterized may fail to provide health benefits. Conversely, granular understanding of conditions essential for lifespan extending paradigms will permit these scenarios to be maximally leveraged to promote health in aging. Accordingly, it is crucial to consider and study the impact of environmental variables to advance studies of aging towards greater rigor and reproducibility. If we are to leverage understanding gained from model systems into improved healthspan and lifespan in humans, a deep understanding of the variables contributing to life trait outcomes is of paramount importance.

## Methods

### Nematode Maintenance

All nematode strains were grown at 20C on nematode growth medium (NGM) agar plates and fed *E. coli* strain OP50-1. NGM agar plates were prepared based on standard protocol [43] using different batches or manufactures of agar. Streptomycin at a final concentration of 0.1875 mg/mL was added in plate preparation to prevent bacteria contamination unless otherwise mentioned. *E. coli* OP50-1 was prepared in LB broth for 48h at 37C incubator without shaking and then kept in 4C unless otherwise noted. OP50 bacteria was seeded onto NGM plates 2 days before use. Other ways of culture OP50 was detailed in Figure legend S1.

The following natural isolates were obtained from the Caenorhabditis Genetics Center (CGC): *C. elegans* N2 Bristol, MY16, JU775; *C. briggsae* AF16, ED3092, HK104; and *C. tropicalis* JU1373. *C. tropicalis* strains JU1630 and QG834 were gifts from Dr. Monica Driscoll’s lab. The listed mutant strains were used: MGH274 *aak-2(ok524) X 4X*, EU31 *skn-1(zu135) IV/nT1 [unc-?(n754) let-?] (IV;V) 3X*, CW152 *gas-1(fc21) X 3X*, MGH433 *raga-1(ok386) II 4X*, MQ887 *isp-1(qm150) IV 3X* were obtained from CGC; SPC227 *skn-1(lax188) IV* was a gift from Dr. Sean Curran’s lab. For aak-2 overexpression, WBM60 *uthIs248 [aak-2p::aak-2(genomic aa1-321)::GFP::unc-54 3’UTR + myo-2p::tdTOMATO]* was obtained from CGC. All experiments were conducted at 20C unless otherwise noted.

### RNA Interference (RNAi)

RNAi clones were isolated from a genome-wide *E. coli* RNAi library, sequence verified, and fed to animals. RNAi feeding plates were prepared using a standard NGM recipe with 5 mM isopropyl-B-D-thiogalactopyranoside and 200 mg/ml carbenicillin. RNAi clones were grown for 18 hours in Luria Broth (LB) containing 200 mg/ml carbenicillin with shaking at 37C. The stationary phase culture was then collected, concentrated through centrifugation, the supernatant was discarded and the pellet was resuspended in LB to 20% of the original culture volume; 250 mL of each RNAi clone concentrate was added to RNAi plates and allowed to dry no more than 48 hours prior to adding the worm embryos or animals.

### Chemicals

Metformin and phenformin were dissolved in water with the stock solutions as 1 M and 0.1 M respectively and were added to OP50 or HT115 bacteria seeded plates 24 hours before adding worms. Paraformaldehyde (PFA) was dissolved in PBS with the stock solutions 4%.

### Longevity Assay

Lifespan analysis was conducted at 20C except where indicated, as previously published [53, 54]. Synchronized L1 animals were seeded onto NGM or RNAi plates and allowed to grow until the L4 stage. On day 0 as indexed in the figure legend, L4 worms were transferred onto NGM or RNAi plates supplemented with 50 mM 5-fluorodeoxyuridine (FUdR) to suppress progeny production. Dead worms were counted every other day. For longevity assay without FUdR, worms were transferred to fresh plates every other day from Day 1 adult to Day 9 adult.

Statistical analysis was performed with online OASIS2 resources [55].

### Paraformaldehyde Treatment of Bacteria

PFA treatment of bacteria was prepared based on a protocol published in 2021 [40]. Briefly, after culturing of OP50, paraformaldehyde at a final concentration of 1% was added to the bacteria, the same volume of PBS was uses as control. PFA or PBS-treated bacteria were shaken at 37 °C in a shaking incubator at 200 rpm for 2 h to allow for even mixing and sufficient exposure to PFA. The treated bacteria were centrifuged at RCF of 3000 × g for 20 min.

Supernatant was removed and 50 mL of LB was added to the pellet to wash off the PFA. The washing was repeated for a total of five times with centrifugation and removal of the supernatant between each washing step. After the final wash, the treated bacteria were resuspended in 5 mL (10-fold concentrated, 10×) to concentrate the aliquots for lifespan assays. The bacteria were seeded on NGM plates and were left to dry for 72 h before the experiments. Bacteria viability was determined by streaking on LB plates to make sure no growth. Metformin or phenformin was added to plates 24 hours before adding worms.

### UV Radiation

After seeding with OP50, dried plates were exposed to UV (9999 × 100 μJ/cm2) for 5 min using a UV Crosslinker (CL-600 Ultraviolet Crosslinker, UVP, USA). Bacteria viability was determined by streaking on LB plates to make sure no growth 3 days after UV-exposure. Metformin or phenformin was added to plates 24 hours before use.

### Quantification and statistical analysis

Statistical analyses were performed using Prism (GraphPad Software). The statistical differences between control and experimental groups were determined by two-tailed Student’s t test (two groups), one-way ANOVA (more than two groups), or two-way ANOVA (two independent experimental variables), with corrected P values < 0.05 considered significant. The log rank test was used to determine significance in lifespan analyses using online OASIS2. Asterisks denote corresponding statistical significance *p < 0.05; **p < 0.01; ***p < 0.001 and ****p < 0.0001.

---

**Figure S1.**
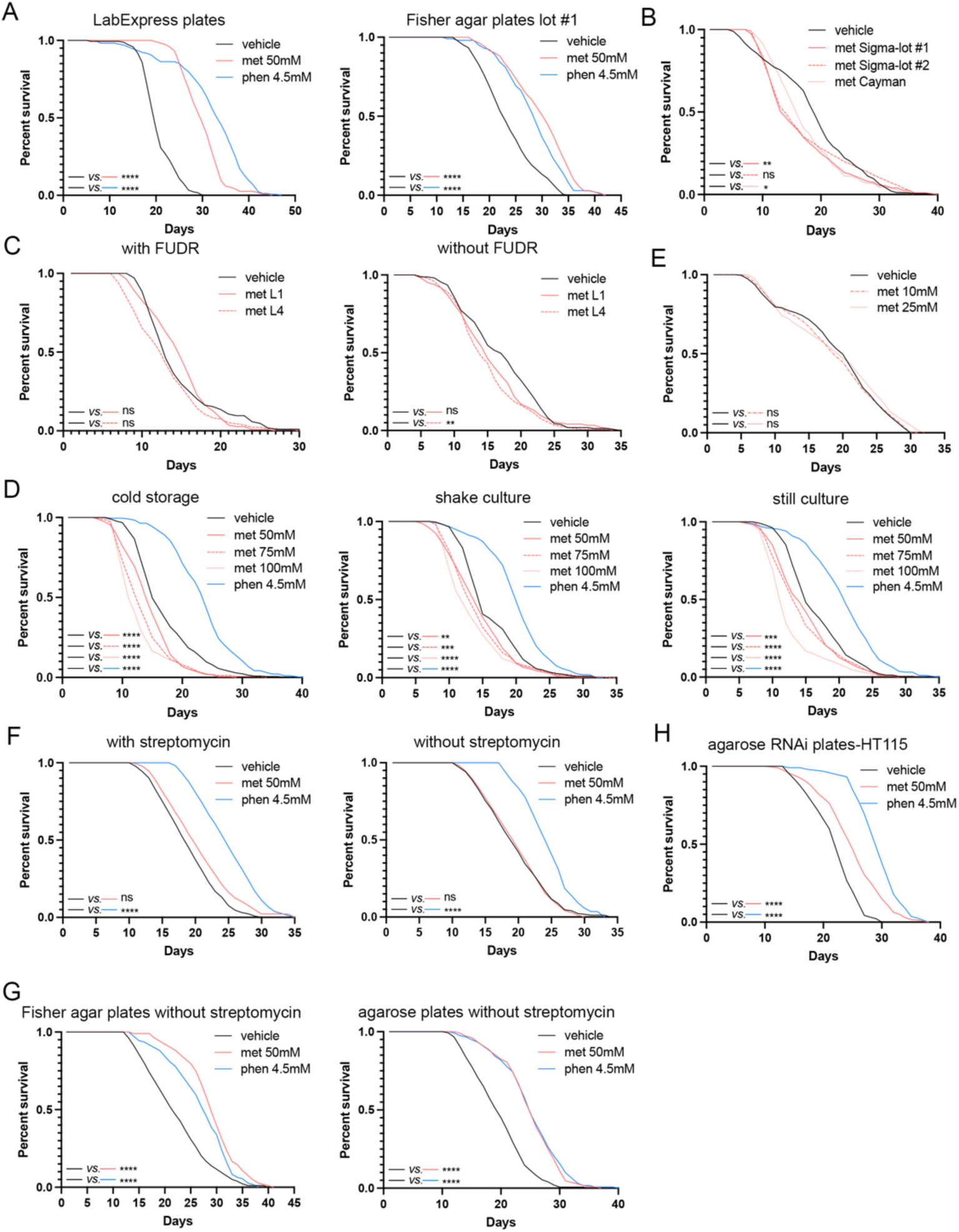
Agar is the only environmental condition that contributes to variation in the lifespan-promoting response to metformin. (A) Metformin and phenformin extend *C. elegans* lifespan in LabExpress and Fisher Scientific agar plates lot #1 (log-rank test). (B) On Bacto agar plates, metformin can not extend lifespan in *C. elegans* using two different lots of metformin obtained from Sigma or one lot from Cayman Chemical (log-rank test). (C) On Bacto agar plates, metformin cannot extend lifespan in *C. elegans*, regardless of whether it is added at the L1 or L4 stage and irrespective of the use of FUdR (log-rank test). (D) On Bacto agar plates, metformin at 50, 75, and 100 mM cannot extend lifespan in *C. elegans*. Phenformin but not metformin is capable of extending lifespan in *C. elegans* feed OP50 bacteria prepared in different ways: cold storage (still culture at 37°C incubator for 48 hours followed by storage in 4°C for 5-7 days before seeding plates), shaken culture (cultured with shaking at 37°C incubator for 18 hours and then immediately seeded onto plates), or still culture (cultured without shaking at 37°C incubator for 48 hours and then immediately seeded) (log-rank test). (E) On Bacto agar plates, metformin at 10 and 25 mM cannot extend lifespan in *C. elegans* (log-rank test). (F) On Bacto agar plates, phenformin but not metformin is able to extend lifespan in *C. elegans* whether or not streptomycin was added during plate preparation (log-rank test). (G) On Fisher agar and agarose plates, metformin and phenformin extend lifespan in *C. elegans* when streptomycin was not added in plate preparation (log-rank test). (H)) On agarose RNAi plates, metformin and phenformin extend lifespan in *C. elegans* when feeding HT115 *E. coli* bacteria (log-rank test). Results are representative of 3–4 biological replicates (see also Table S1 for tabular results and biological replicates). *p < 0.05, **p < 0.01, ***p < 0.001, ****p < 0.0001.

**Figure S2.**
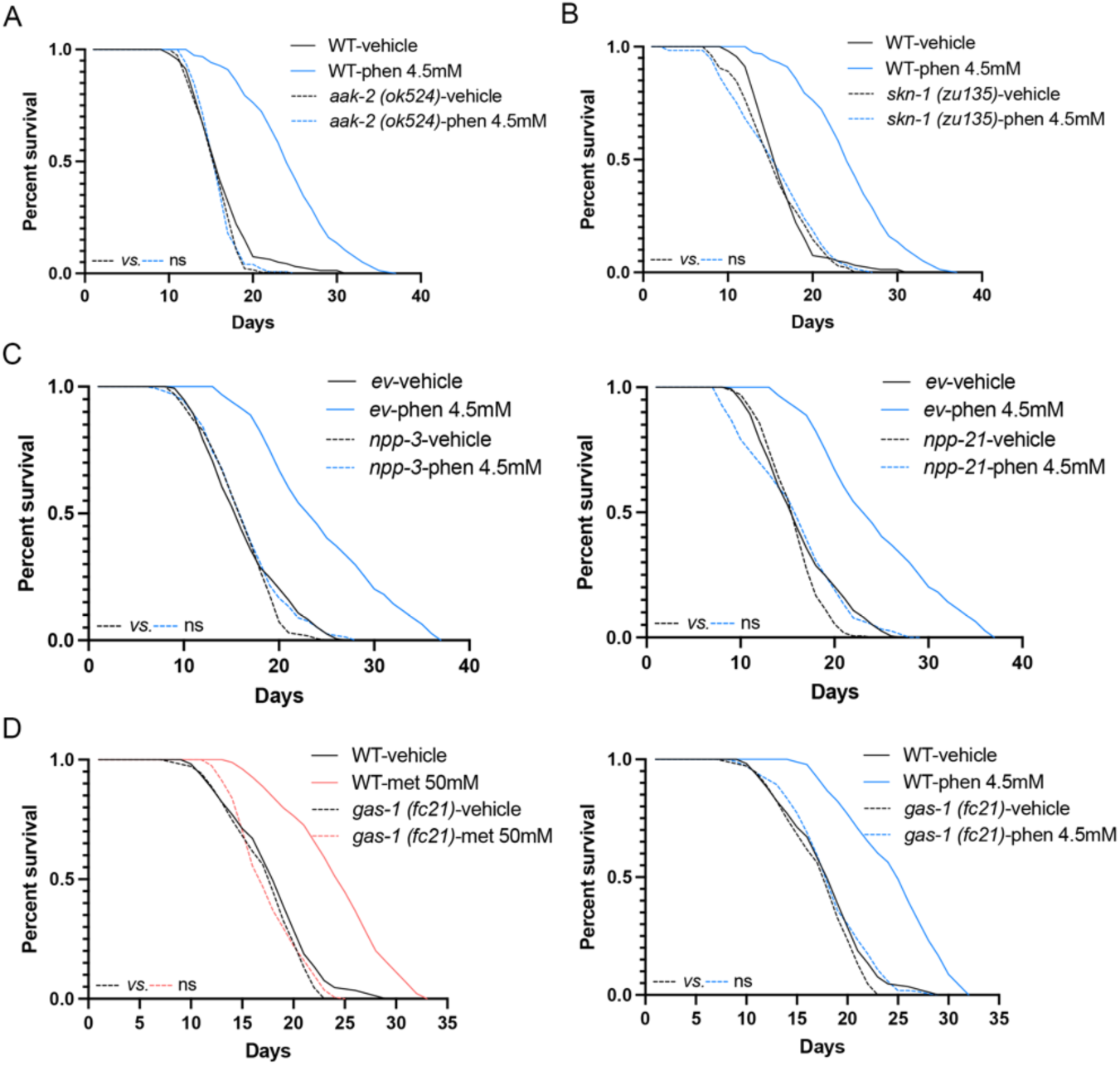
Metformin and phenformin share the same genetic dependency. (A-C) Phenformin cannot extend *C. elegans* lifespan in *aak-2 (ok524)* and *skn-1 (zu135)* mutants, or feeding *npp-3, npp-21* RNAi bacteria (log-rank test). (D) On agarose plates, metformin or phenformin could not extend *C. elegans* lifespan in *gas-1 (fc21)* mutants (log-rank test). Results are representative of 3–5 biological replicates (see also Table S2).

**Figure S3.**
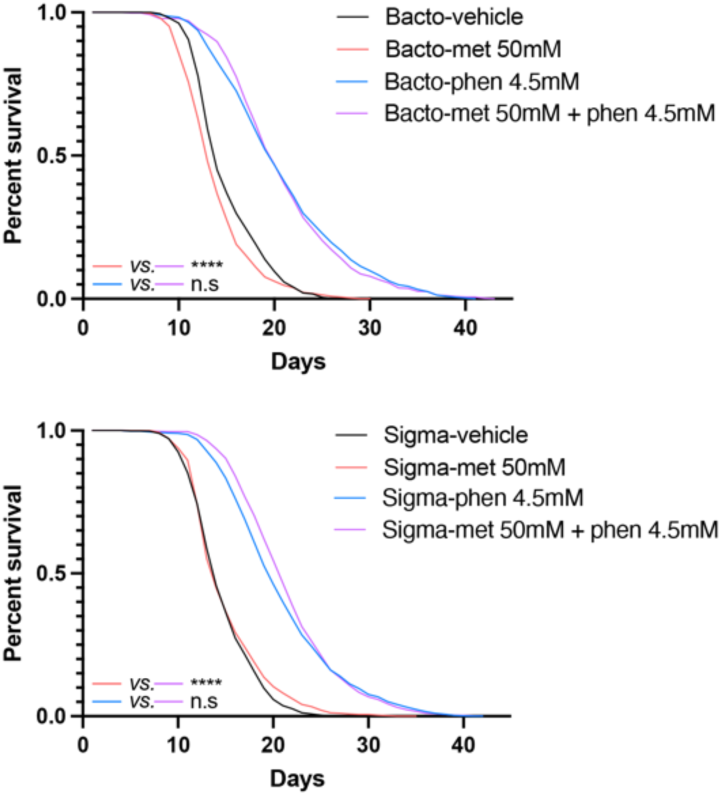
Metformin and phenformin have no additivity in lifespan when using Bacto or Sigma agar plates. When using Bacto or Sigma agar plates, the addition of metformin to phenformin does not result in any additional increase in the lifespan of *C. elegans* compared to treatment with phenformin alone (log-rank test). Results are representative of 4–5 biological replicates (see also Table S2 for tabular data and biological replicates). ****p < 0.0001.

**Figure S4.**
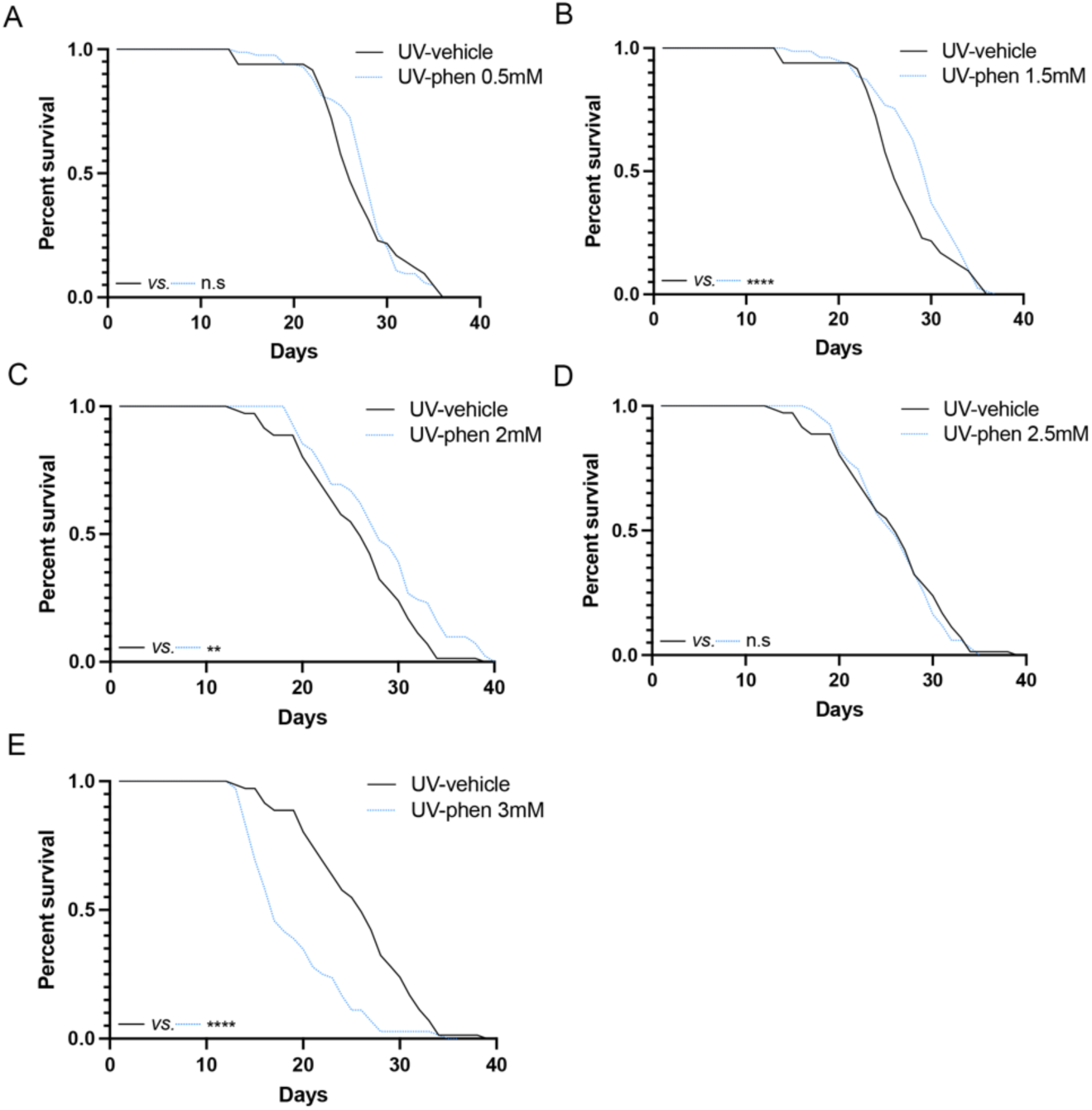
Phenformin extends *C. elegans* lifespan when fed UV-killed bacteria. On agarose plates containing UV-killed bacteria, compared to vehicle, phenformin at 0.5 mM (A) and 2.5 mM (D) does not have any significant impact in lifespan, whereas phenformin at 1.5 mM (B) and 2 mM (C) extend the lifespan of *C. elegans*; phenformin at 3 mM (E) results in a shortened lifespan (log-rank test). Results are representative of 4 biological replicates (see also Table S3 for tabular data and biological replicates). **p < 0.01, ****p < 0.0001.

